# Differences between Scaly-sided Merganser (*Mergus squamatus*) and Common Merganser (*M. merganser*) feather microstructure

**DOI:** 10.1101/2025.01.30.635631

**Authors:** Donghong Li, Shiyu Zhang, Ran Tian, Yongbin Zhao, Guodong Yi

## Abstract

The microstructures of breast feathers, flank feathers, rectrices, and down feathers of both Scaly-sided Merganser (*Mergus squamatus*) and Common Merganser (*M. merganser*) were meticulously examined using scanning electron microscope. The observed data were subjected to the Mann-Whitney *U* test, followed by hierarchical clustering and Fisher’s discriminant analysis of indicators with significant differences in morphological structure. The findings revealed statistically significant differences between Scaly-sided Merganser and Common Merganser in terms of the length of the base handle from distal barbules in the breast feathers, the length of the base handle, the number of hooked barbules, the number of cilia from distal barbules, and the number of ventral teeth from proximal barbules in rectrices. Additionally, variations in the distance between nodules, the nodular diameter, and the length of the base nodular of the down feathers were noted. Hierarchical clustering successfully categorized Scaly-sided Merganser and Common Merganser into distinct groups, achieving a 100% accuracy rate. Moreover, Fisher’s discriminant analysis demonstrated a 100% correctness rate when the back-substitution method was used to test the discriminant equation. Consequently, the length of the base handle of distal barbules in breast feathers, along with all selected indicators from rectrices and down feathers, may provide references to identify Scaly-sided Merganser and Common Merganser.

## Introduction

The microstructure of bird feathers exhibits significant variation among species while maintaining strong stability and interspecific specificity[1]. Chandler first proposed feathers as a core and distinctive external feature of birds, suggesting that their microscopic structure, especially that of barbules, is potentially taxonomically significant and diagnostically indicative of bird species[2]. A notable example is Roxie, who confirmed that the common starling (*Sturnus vulgaris*) was the cause of an aviation accident by analyzing feathers found in the airplane wreckage[3]. To date, many research findings have demonstrated the significant value of feather microstructural features in the identification and classification of bird species, providing crucial evidence for avian taxonomy research[4–8].

Scaly-sided Merganser (*Mergus squamatus*) has been listed as endangered by the International Union for Conservation of Nature (IUCN) since 2002[9]. Consequently, there is substantial concern regarding various aspects of Scaly-sided Merganser, including population size[10–12], habitat selection[13,14], genetic diversity[15], migration connections[16], and breeding ecology[17]. In addition to field observations, molecular biology and isotope tracing are often used to address these issues. However, due to the rarity and high vigilance of Scaly-sided Mergansers, direct capture of blood or tissue samples is becoming increasingly challenging. Thus, naturally shed feathers might serve as effective research material. It is noteworthy that Common Merganser (*M. merganser*), classified as the species of least concern according to the IUCN Red List, is distinguished as the largest and most widely distributed species within the genus *Mergus* in China[18]. Both species belong to the order Anseriformes, family Anatidae, and genus *Mergus*, which share similar ecological behaviors and exhibit some degree of habitat overlap[19]. This poses a challenge for ecologists, as feather samples collected may include contributions from both species. Nonetheless, when conducting molecular biology studies on feathers or performing isotope analysis, it is imperative to meticulously differentiate the feathers of Scaly-sided Merganser from those of other species. The uncontrollable timing of feather collection often leads to significant DNA degradation after sample acquisition. Additionally, naturally shed feathers inherently contain a limited amount of DNA, posing considerable challenges for molecular biology analysis. Therefore, morphological differentiation of feather samples is imperative. In the case of both Scaly-sided Merganser and Common Merganser, the macroscopic morphology of their feathers is remarkably similar, making distinguishing between the two species challenging through simple observation alone. Hence, a detailed investigation of the microstructure is necessary for precise identification.

This study aimed to investigate the microstructural characteristics of the breast feathers, flank feathers, rectrices, and down feathers of both Scaly-sided Merganser and Common Merganser. Quantitative analysis of these features will be conducted to identify differences between them, with the goal of providing a basis for their recognition.

## Materials and methods

### Materials and processing

Feathers from the Scaly-sided Merganser were gathered during their winter molt at Longhu Mountain, Jiangxi Province, China, involving approximately 20 individuals. These feathers were then classified according to Featherbase[20]. In addition, feathers of Common Merganser were obtained from an adult female at the Animal Specimen Museum, Jilin Normal University, Siping City, Jilin Province, China. For each species, five feathers were selected from each of breast feathers, flank feathers, rectrices, and abdominal down feathers. The research area was defined as the middle part of the feather vanes. Considering that down feathers do not form vanes, nodular barbules were extracted from the middle part as the research region. After cleaning with tap water and 75% alcohol, we air-dried the feathers before carefully detaching the barbules from the rachises and affixing them to conductive adhesive tape. Images and data regarding various variables such as the length of the base handle, the number of hooked barbules, the number of cilia, the number of ventral teeth, the distance between nodules, the nodular diameter, and the length of the base nodular were captured and recorded using a Hitachi TM4000 scanning electron microscope (Hitachi, Tokyo, Japan). 30 datasets were collected for each variable of the two species.

### Statistical analyses

Statistical analyses were conducted using SPSS 25.0 software (IBM Corporation, Armonk, NY, USA). Given the small sample sizes, the data were analyzed using the Mann-Whitney *U* test[4]. The results are presented in the form of median (interquartile range). Differences at *p* < 0.05 were considered significant. For morphological structures exhibiting significant differences, hierarchical clustering and Fisher’s discriminant analysis were performed. The morphological indicators displaying significant differences served as clustering factors for the cluster analysis, with the number of clustering groups set at 2. Fisher’s discriminant functions were formulated, and the variance contribution rate and the cumulative contribution rate of each function were computed. The discriminant ability was tested for significance using Wilks’ lambda, where *p* < 0.05 indicated significant discriminant ability. The accuracy of the discriminant functions was validated through the back-substitution method[21].

## Results

### Microstructure of breast feathers, flank feathers, rectrices, and down feathers

The breast feathers, flank feathers, and rectrices of Scaly-sided Merganser and Common Merganser are predominantly composed of distal and proximal barbules. The distal barbules consisted of the base and the pennulum, bearing both hooked barbules and cilia. In contrast, the proximal barbules feature ventral teeth and cilia. The down feathers consisted of nodular barbules.

The proximal barbules of the breast feathers in Scaly-sided Merganser are parallel and relatively sparse, featuring an enlarged base with a flattened shape. There are 2-3 triangular ventral teeth. Cilia grow unilaterally, numbering 4-6 (Fig 1a). The distal barbules are similar in length to the proximal barbules. Hooked barbules, usually 4, are noticeably curved and directed toward the base. Cilia are unilateral, 5-7, and gradually increase in length from the tip of the pennulum to the base (Fig 1b). Common Merganser and Scaly-sided Merganser have similar microstructures in their breast feathers. The proximal barbules have a flattened basal part and run parallel to each other. Typically, the ventral teeth are 2-4 in number. A few short cilia were present at the tip of the pendulum (Fig 1c). The distal barbules displayed basal expansion, with unilaterally growing hooked barbules numbering 4-5 and presenting a blunt anterior aspect. The tip of the pennulum exhibited cilia, typically consisting of 5-6 solitary cilia (Fig 1d).

**Fig 1.**
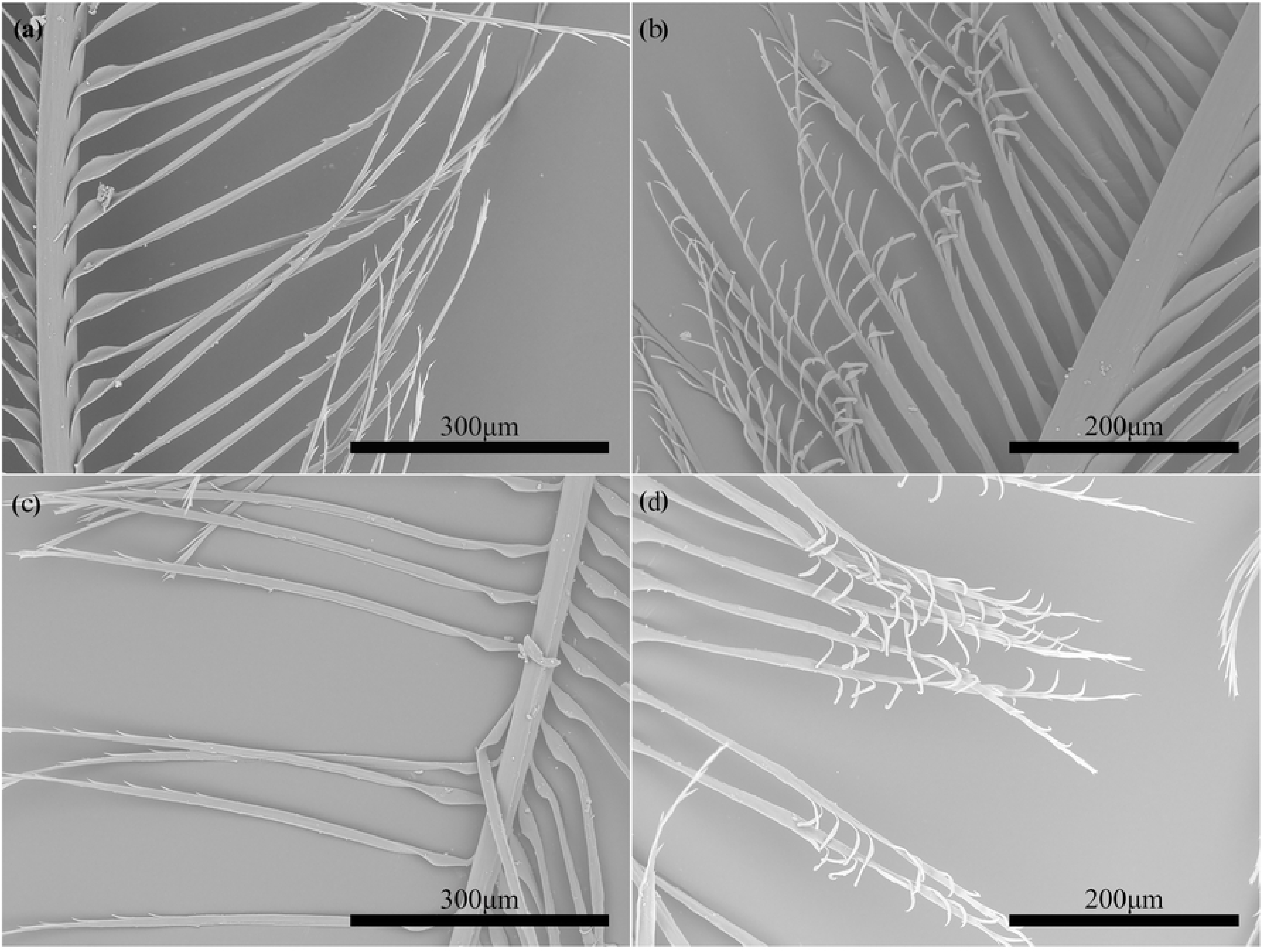
The microstructure of breast feathers in Scaly-sided Merganser and Common Merganser. (a) The proximal barbules of breast feathers in Scaly-sided Merganser. (b) The distal barbules of breast feathers in Scaly-sided Merganser. (c) The proximal barbules of breast feathers in Common Merganser. (d) The distal barbules of breast feathers in Common Merganser.

The proximal barbules of the flank feathers in Scaly-sided Merganser are arranged on the same side of the branches in a parallel distribution. The basal stipe is flattened, and the ventral teeth exhibit a serrate pattern, typically numbering 2-3 (Fig 2a). The distal barbules were arranged parallel to each other, featuring 3-4 hooked barbules. Cilia were observed at the pendulum, exhibiting a progressive reduction in length. They were unilaterally present and typically numbered 6-7 (Fig 2b). Common Merganser and Scaly-sided Merganser have similar microstructures in their flank feathers. The ventral teeth of the proximal barbules were arranged in a trapezoidal pattern, featuring 2-3 teeth (Fig 2c). In contrast, the distal barbules possessed 4-5 hooked barbules and 5-8 solitary cilia (Fig 2d).

**Fig 2.**
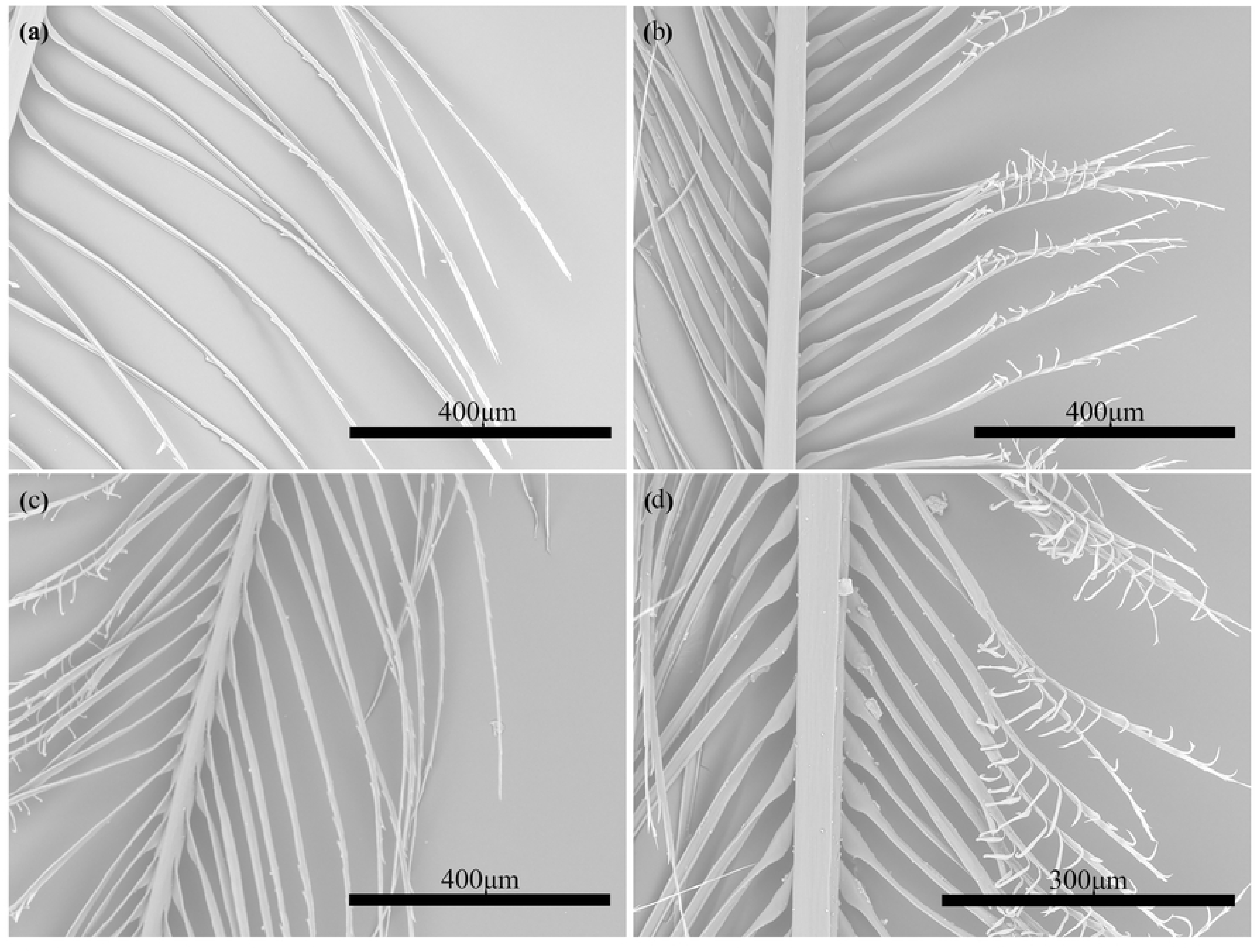
The microstructure of flank feathers in Scaly-sided Merganser and Common Merganser. (a) The proximal barbules of flank feathers in Scaly-sided Merganser. (b) The distal barbules of flank feathers in Scaly-sided Merganser. (c) The proximal barbules of flank feathers in Common Merganser. (d) The distal barbules of flank feathers in Common Merganser.

The proximal barbules of the rectrices in Scaly-sided Merganser are densely arranged on one side, with 4-5 serrated ventral teeth, while the tips of the proximal barbules taper and extend, forming a filamentous structure (Fig 3a). In contrast, the distal barbules, which are shorter than their proximal counterparts, align in parallel with close spacing, featuring long and curved hooked barbules typically numbering 5-7 in the middle. The cilia exhibit a filiform nature and are positioned opposite each other, with one long and the other short, typically forming 6-8 pairs, while the terminal barbules undergo differentiation, assuming an elongated and filamentous structure (Fig 3b). The rectrices of Common Merganser and Scaly-sided Merganser exhibit identical barbule structures. The ends of proximal barbule processes differentiate into long filamentous structures. The ventral teeth are serrated, typically numbering 5-6 (Fig 3c). The distal barbules, which are shorter than their proximal counterparts, are closely arranged and adorned with 6-8 hooked barbules. The cilia form 7-11 pairs, exhibiting a minor discrepancy in length, and the ends of the barbules differentiate into long filamentous structures (Fig 3d).

**Fig 3.**
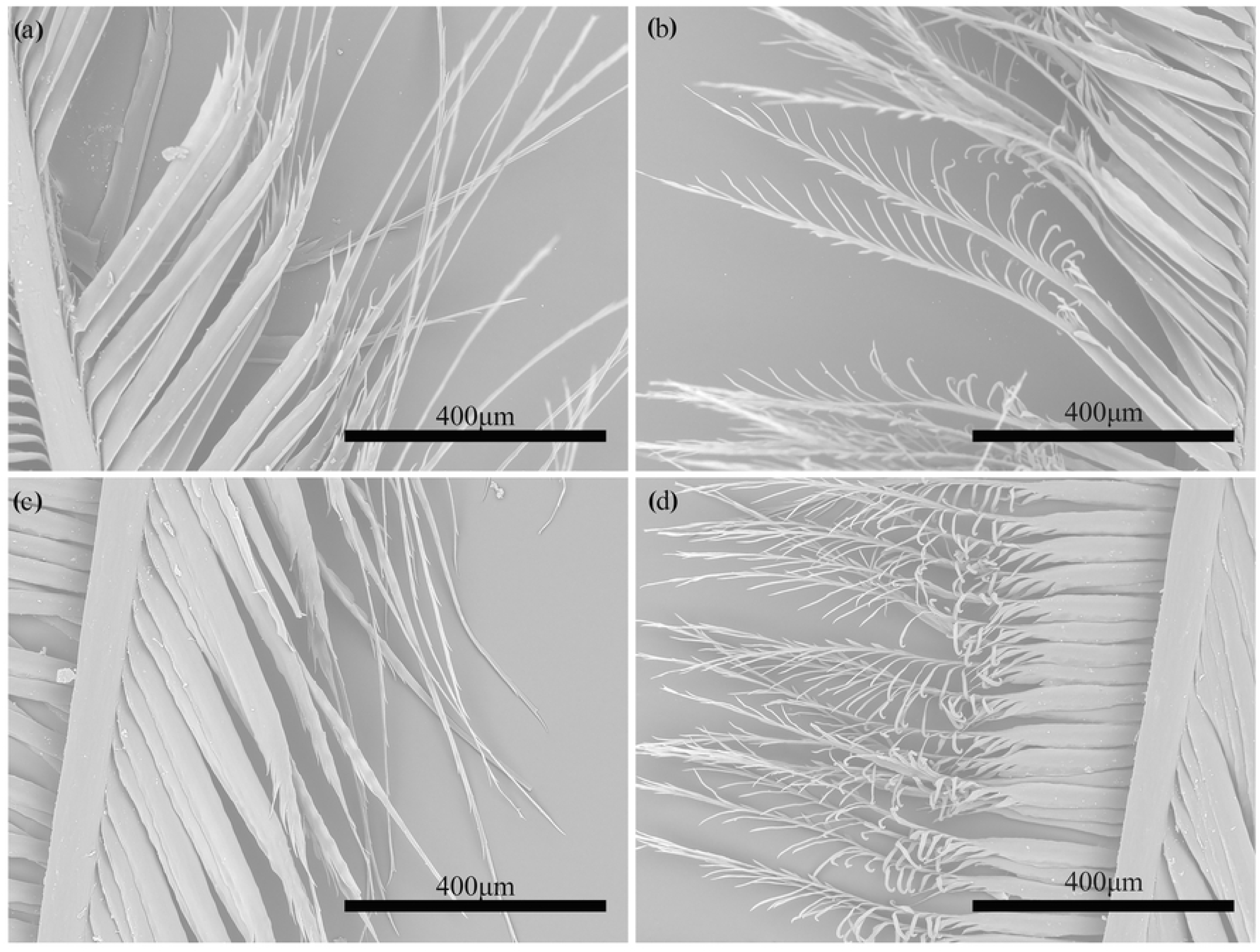
The microstructure of rectrices feathers in Scaly-sided Merganser and Common Merganser. (a) The proximal barbules of rectrices in Scaly-sided Merganser. (b) The distal barbules of rectrices in Scaly-sided Merganser. (c) The proximal barbules of rectrices in Common Merganser. (d) The distal barbules of rectrices in Common Merganser.

The down feathers of Scaly-sided Merganser display nodular barbules that are loosely arranged along the middle and upper sections of the feather shaft, characterized by their slender morphology, with each nodular barbule comprising several inflated triangular nodes, numbering 3-4, with irregular spacing between these nodes (Fig 4a). The microstructure of the down feathers of Common Merganser and Scaly-sided Merganser displayed minimal differences, with the nodular barbules of Common Merganser appearing filamentous and shorter. They typically terminated with 1-3 triangular nodes at their ends, exhibiting irregular spacing (Fig 4b).

**Fig 4.**
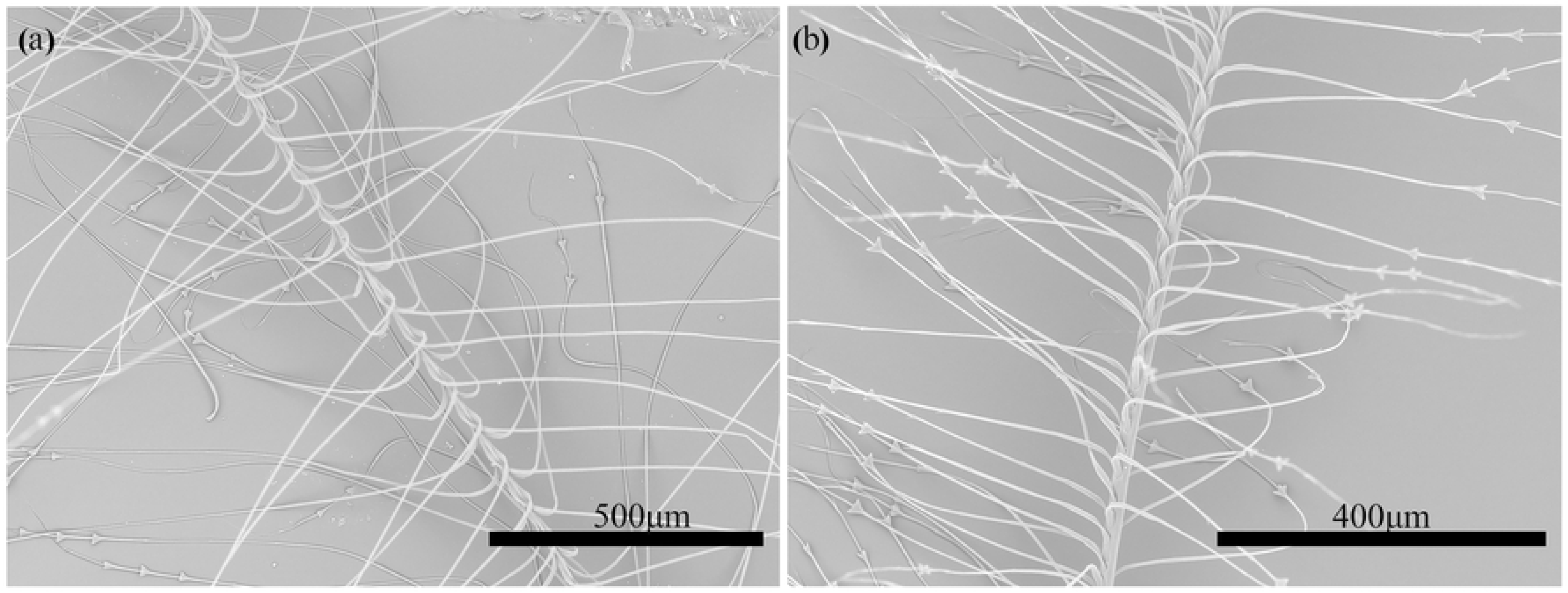
The microstructure of down feathers in Scaly-sided Merganser and Common Merganser. (a) Scaly-sided Merganser. (b) Common Merganser.

### Analysis of the differences in feather microstructure

Scaly-sided Merganser and Common Merganser exhibit significant differences in multiple microstructural feather features (Table 1). Notably, variations were observed in the length of the base handle from distal barbules in breast feathers. These differences extended to the length of the base handle, the number of hooked barbules, cilia from distal barbules, and ventral teeth from proximal barbules in rectrices. Additionally, differences in the distance between nodules, the nodular diameter, and the length of the base nodular were noted in down feathers. However, no statistically significant differences were found in the number of hooked barbules, cilia from distal barbules, or ventral teeth from proximal barbules in the breast feathers between Scaly-sided Merganser and Common Merganser. Similarly, there were no significant variations in the length of the base handle, the number of hooked barbules, cilia from distal barbules, or the number of ventral teeth from proximal barbules in the flank feathers. In summary, the length of the base handle from distal barbules in breast feathers, in conjunction with selected metrics of rectrices and down feathers, differed significantly between Scaly-sided Merganser and Common Merganser.

**Table 1.**
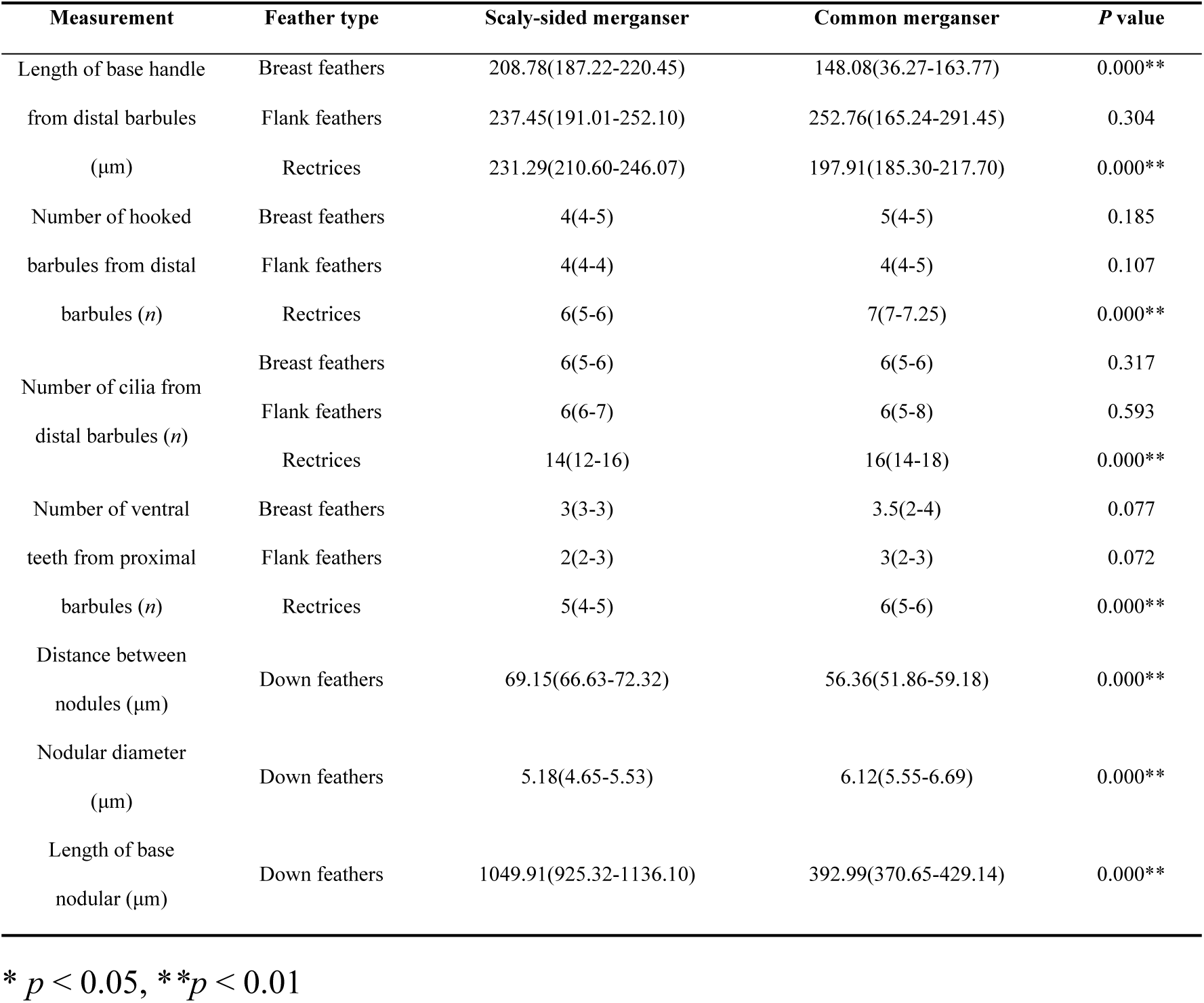
Comparison of the barbule structure between Scaly-sided Merganser and Common Merganser.

### Hierarchical clustering of characterization indicators of feathers

Matrix scatter plots were generated using key indicators of significant differences in feather morphology and structure. These indicators included the length of the base handle from distal barbules in breast feathers, the length of the base handle, the number of hooked barbules, the number of cilia from distal barbules, and the number of ventral teeth from proximal barbules in rectrices, as well as the distance between nodules, the nodular diameter, and the length of the base nodular in down feathers. The plots distinctly illustrate the separate clustering of Scaly-sided Merganser and Common Merganser (Fig 5a). These significant indicators were subsequently employed as variables for hierarchical clustering and cluster purity calculations. The results demonstrated that Scaly-sided Merganser formed one distinct category, while the common mergers formed another (Fig 5b), achieving a 100% accuracy rate.

**Fig 5.**
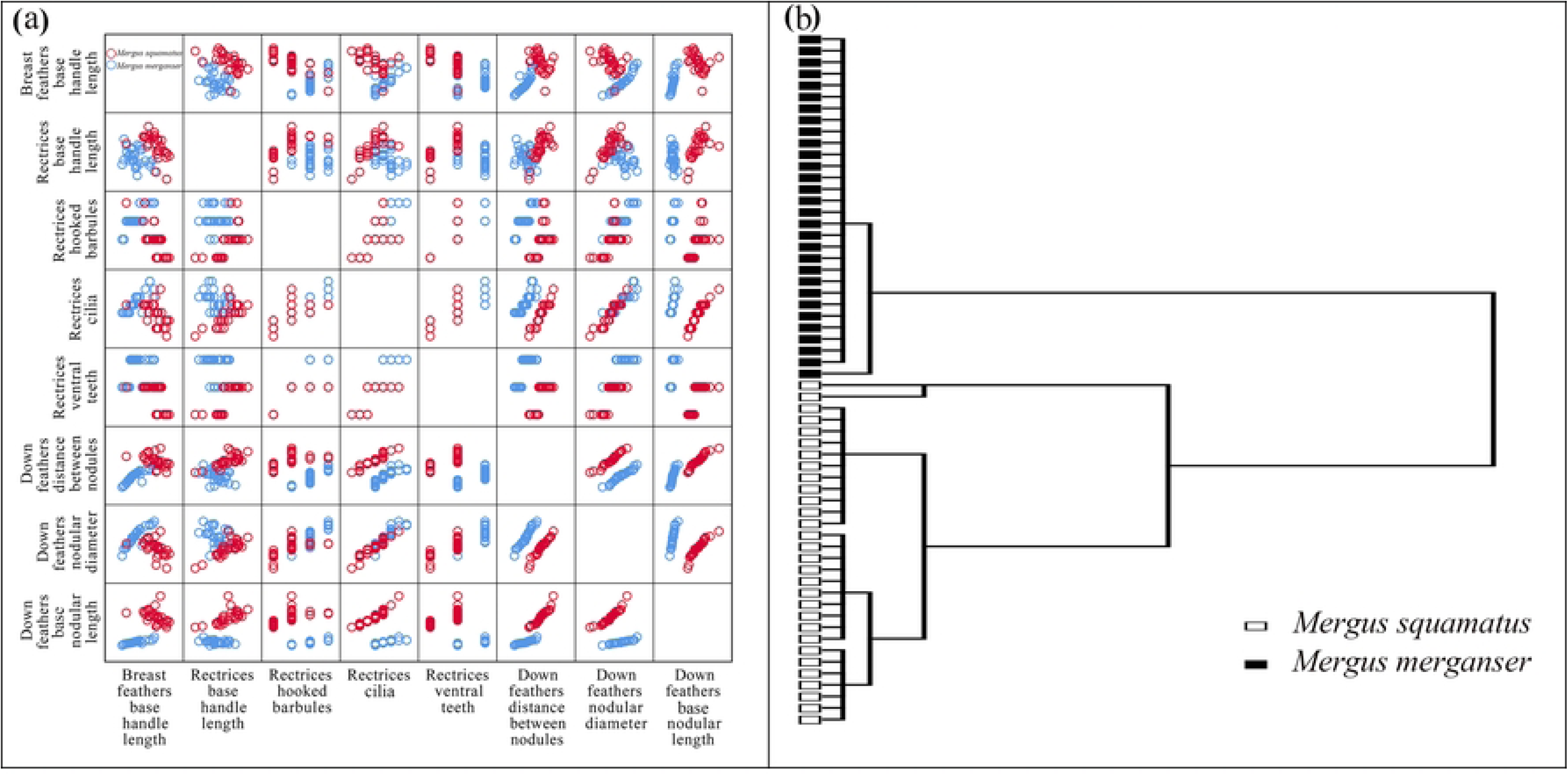
Cluster results diagram. (a) Matrix scatterplot based on the length of the base handle from distal barbules in breast feathers, the length of the base handle, the number of hooked barbules, the number of cilia from distal barbules, and the number of ventral teeth from proximal barbules in rectrices, as well as the distance between nodules, the nodular diameter, and the length of the base nodular in down feathers. (b) Hierarchical clustering based on the length of the base handle from distal barbules in breast feathers, the length of the base handle, the number of hooked barbules, the number of cilia from distal barbules, and the number of ventral teeth from proximal barbules in rectrices, as well as the distance between nodules, the nodular diameter, and the length of the base nodular in down feathers.

### Fisher’s discriminant analysis of feather characterization indices

Fisher’s discriminant analysis utilized indicators reflecting significant differences in feather morphological structure as variables, resulting in a distinct discriminant function denoted as *Y* (Table 2). The accuracy of the discriminant equation was assessed through variance contribution following the principles of Fisher’s discriminant analysis. The findings reveal that the cumulative variance contribution of *Y* reaches 100%, offering a comprehensive representation of the sample information (Table 3). Consequently, the subsequent typical discriminant function is derived:

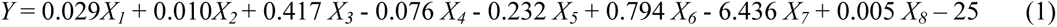

**Table 2.**
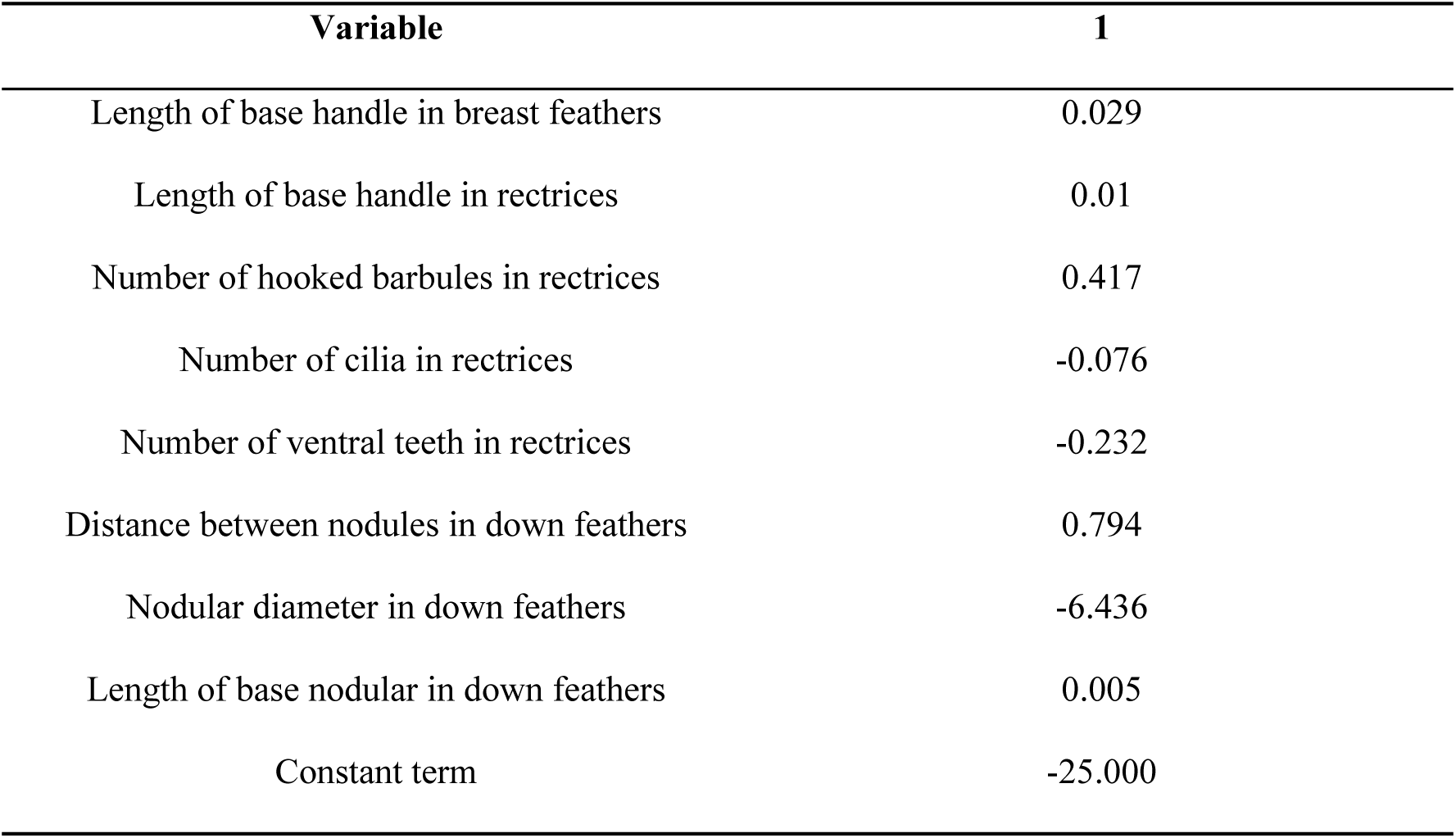
Coefficient and constant terms of the typical discriminant function.

**Table 3.** Variance in the discriminant function of feathers and its significance.

| Function | Eigenvalue | Variance<br>contribution (%) | Cumulative variance<br>contribution (%) | Typical relevant<br>coefficients |
| --- | --- | --- | --- | --- |
| 1 | 132.296 | 100 | 100 | 0.996 |

In Equation (1), the variables *X_1_* to *X_8_* represent specific features related to the morphological structure of feathers: the length of the base handle from distal barbules in breast feathers, the length of the base handle, the number of hooked barbules, the number of cilia from distal barbules, the number of ventral teeth from proximal barbules in rectrices, the distance between nodules, the nodular diameter, and the length of the base nodular in down feathers. Collectively, these characteristics serve as indicators for assessing feather morphology. By employing the *Wilks’ Lambda test*, it was observed that the function *Y* had a significant (*p* < 0.05) ability to discriminate between the species of Scaly-sided Merganser and Common Merganser when indicators with significant differences in feather morphology and structure were used as variables. After a thorough examination of the data used to construct the discriminant function on a case-by-case basis using the back-substitution method, the accuracy of the discriminant equation obtained was 100%.

## Discussion

Previous studies have demonstrated that feather microstructures exhibit variations not only among different species but also between sexes within the same species[4–6]. In this study, we observed and measured breast feathers, flank feathers, rectrices, and down feathers of Scaly-sided Merganser and Common Merganser, followed by rigorous statistical analyses. The results revealed significant differences in certain characteristics between the two species, including the length of the base handle from distal barbules in breast feathers, the length of the base handle, the number of hooked barbules, the number of cilia from distal barbules, and the number of ventral teeth from proximal barbules in rectrices, as well as the distance between nodules, the nodular diameter, and the length of the base nodular in down feathers. Consequently, these distinctions could potentially serve as a basis for differentiating between the Scaly-sided Merganser and the Common Merganser.

The microstructure of feathers provides an important reference for the identification of bird species[1]. However, species identification cannot be achieved using a single indicator alone; it requires the evaluation of multiple features. In this study, hierarchical clustering and Fisher’s discriminant analysis were applied to morphological traits that exhibited significant differences. Hierarchical clustering successfully separated the two species into distinct branches with 100% accuracy, while the Fisher discriminant function also achieved 100% precision. These results suggest that incorporating multiple indicators in the analysis can significantly enhance the accuracy and reliability of species identification.

This study on the microstructural differences in the feathers of the Scaly-sided Merganser and Common Merganser may provide a basis for their identification. It also aids in the further exploration of their ecological characteristics, including habitat utilization, dietary preferences, and breeding behaviors. However, several limitations merit consideration. First, our study was limited by the limited number of feather samples, which hindered sample diversity and representativeness, potentially introducing errors into the results. Second, the absence of samples of remiges from Scaly-sided Merganser limits our understanding of the differences between the two species. Therefore, in future research, further verification and the establishment of a database will become key research goals. It is recommended to increase the sample size and consider various factors, including the influence of body size, flight ability, gender, and seasonal changes on the microstructure of bird feathers. At the same time, it is suggested to integrate morphological and molecular biological methods to further improve the accuracy of classification.

## Conclusion

Our study successfully identified significant microstructural differences in the feathers of Scaly-sided Merganser and Common Merganser, demonstrating the potential of feather morphology as a reliable method for species differentiation. The findings underscore the necessity of utilizing multiple morphological indicators for accurate identification, as evidenced by the 100% accuracy achieved through hierarchical clustering and Fisher’s discriminant analysis. This research not only contributes to the understanding of avian taxonomy but also provides a foundation for further ecological studies regarding habitat use and behavioral patterns in these species. Future investigations should aim to expand sample sizes and incorporate additional factors to enhance the robustness of feather microstructure analysis in avian classification.

## Acknowledgments

This work was financially supported by the Jilin Provincial Department of Science and Technology (grant number: 20220101331JC). We thank AJE for assistance with the revision of the paper.

## Author Contributions

**Conceptualization:** Donghong Li, Shiyu Zhang.

**Formal analysis:** Donghong Li, Yongbin Zhao.

**Funding acquisition:** Yongbin Zhao.

**Investigation:** Donghong Li.

**Methodology:** Donghong Li, Shiyu Zhang.

**Resources:** Guodong Yi.

**Supervision:** Shiyu Zhang, Ran Tian, Yongbin Zhao.

**Visualization:** Donghong Li, Yongbin Zhao.

**Writing– original draft:** Donghong Li.

**Writing– review & editing:** Shiyu Zhang, Ran Tian, Yongbin Zhao, Guodong Yi

